# Methanotroph phenotypic heterogeneity in a methane-oxygen counter gradient

**DOI:** 10.1101/2023.10.05.561118

**Authors:** Delaney G. Beals, Aaron W. Puri

## Abstract

Connecting genes to phenotypic traits in bacteria is often challenging because of a lack of environmental context in laboratory settings. Laboratory-based model ecosystems offer a means to better account for environmental conditions compared to standard planktonic cultures, and can help link genotypes and phenotypes. Here, we present a simple, cost-effective, laboratory-based model ecosystem to study aerobic methane-oxidizing bacteria (methanotrophs) within the methane-oxygen counter gradient typically found in the natural environment of these organisms. Culturing the methanotroph *Methylomonas* sp. strain LW13 in this system resulted in formation of a distinct horizontal band at the intersection of the counter gradient, which we discovered was not due to increased numbers of viable bacteria at this location but instead to an increased amount of polysaccharides. We also discovered that different methanotrophic taxa form polysaccharide bands with distinct locations and morphologies when grown in the methane-oxygen counter gradient. By comparing transcriptomic data from LW13 growing within and surrounding this band, we identified genes upregulated within the band and validated their involvement in growth and band formation within the model ecosystem using knockout strains. Notably, deletion of these genes did not negatively affect growth using standard laboratory conditions. This work highlights the use of a laboratory-based model ecosystem that more closely mimics the natural environment to uncover methanotroph phenotypes missing from standard planktonic cultures, and to link these phenotypes with their genetic determinants.

## INTRODUCTION

Despite the explosion in availability of bacterial genomic data, linking genes with organismal phenotypes remains difficult. One reason is that assigning gene functions can be hindered by a lack of ecological context in the lab, as bacteria are removed from the environments where they evolved. Laboratory-based model ecosystems are a powerful tool that can be used to more closely recapitulate the natural environment than simple planktonic cultures [1, 2].

Aerobic methane-oxidizing bacteria (methanotrophs) play important roles in biogeochemical cycling, bioremediation, and sequestration of the potent greenhouse gas methane [3–5]. These bacteria require methane as a carbon and energy source, and molecular oxygen as an electron acceptor and atom donor in the methane monooxygenase reaction. In the environment, methanotrophs often exist in opposing gradients of oxygen and methane, where oxygen diffuses from the atmosphere or water column above, and methane diffuses up from anaerobic decomposition processes below [6–9]. Studies using methane enrichments of sediment have consistently observed dynamic population flux amongst methanotrophs in response to shifting oxygen and methane availability [10, 11]. Methanotrophs are therefore particularly shaped by the spatial heterogeneity of their environment, however in the laboratory these organisms have primarily been studied using homogenous planktonic cultures. This is likely impeding our ability to both understand the molecular details of methanotroph physiology, and to optimize these details for industrial processes and climate change mitigation [12, 13].

Culturing methanotrophs within a spatially resolved model ecosystem may help uncover phenotypes that are otherwise undetectable in planktonic cultures. Previously, researchers demonstrated that growing methanotrophs within opposing gradients of methane and oxygen in the lab is feasible using a variety of methods. In some examples, soil was suspended on a synthetic polymer membrane that allowed gas exchange from the methane supply below [6, 14, 15]. In other cases, sediment was added to semi-solid agarose and exposed to opposing chambers of air and methane [16, 17]. To our knowledge, these systems have been primarily used to isolate and classify methanotrophs from mixed methane-oxidizing consortia and have not yet been used to collect molecular-level information about the physiology of methanotrophs under opposing methane and oxygen gradients.

Inspired by the methane-taxis assay used by Dedysh and coworkers [18], we created a simple, inexpensive, laboratory-based model ecosystem using a disposable syringe that enables researchers to culture methanotrophs in a spatially resolved context. We first characterized the counter gradients of methane and oxygen in this model ecosystem, referred to as the gradient syringe, and then cultured individual methanotrophs within the system. Using this method, we observed a horizontal band form at the intersection of the counter gradient even though the entire system was uniformly inoculated with bacterial cells. We discovered that this band was due to an increased amount of polysaccharides at this depth, but not an increased number of viable cells. Through transcriptional analysis across layers of the gradient syringe, we identified many genes with differential expression in *Methylomonas* sp. strain LW13 (LW13) growing at different depths of the system. We subsequently confirmed using genetics that three of these genes are important for production of the polysaccharide band, and that these genes are also required for growth in the gradient syringe but not in planktonic culture. These results provide insights into the heterogenous physiological responses of methanotrophs growing under the opposing methane and oxygen gradients they encounter in the environment and highlight the potential of this laboratory-based model ecosystem to uncover the genes responsible for these phenotypes.

## RESULTS

### Different methanotrophic taxa form horizontal bands with distinct locations and morphologies when grown in the gradient syringe model ecosystem

We established a simple laboratory-based model ecosystem to cultivate methanotrophs using semi-solid agarose in disposable syringes (**Fig 1A**). Syringes were equipped with a sterile filter tip and filled with agarose, bacterial inoculum, and nitrate mineral salt medium. We inoculated gradient syringes with individual cultures of four methanotrophs isolated from Lake Washington, Seattle, USA [19]. All methanotrophs inoculated into the gradient syringe formed a horizontal band around the midpoint of the agarose plug in as few as three days, even though the agarose was uniformly inoculated with bacteria, a phenomenon previously observed in more complex setups [16, 17]. With additional days of incubation and replenishment of methane in the syringe headspace, this band became more pronounced and opaque.

**FIGURE 1.**
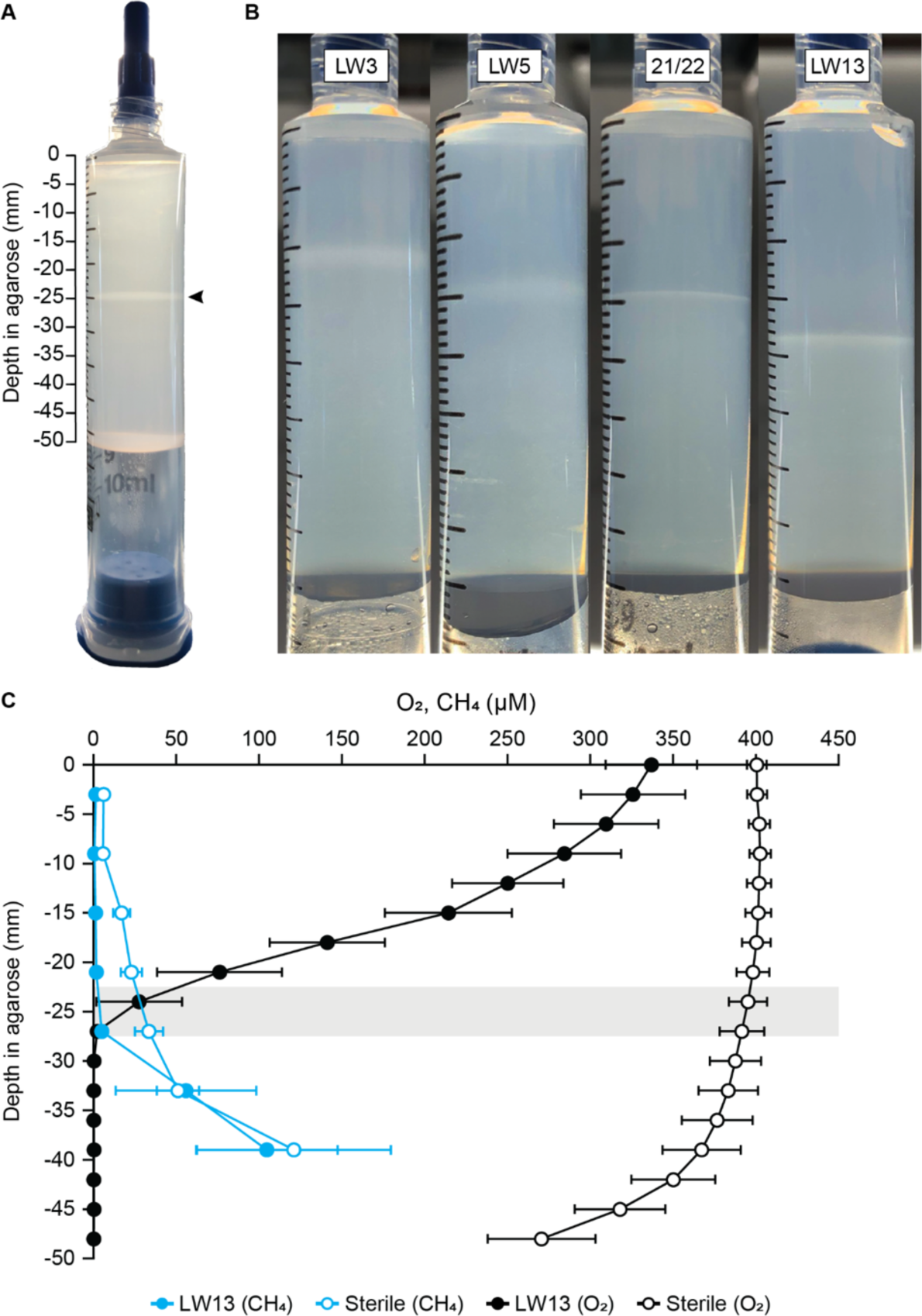
(**A)** The gradient syringe model ecosystem inoculated with the methanotroph *Methylomonas* sp. strain LW13. Black arrowhead indicates location of the horizontal band. (**B)** Cultivation of the Lake Washington-isolated methanotrophic strains in the gradient syringe, *Methylosinus* sp. strain LW3 (day 7 within the syringe), *Methylobacter tundripaludum* 21/22 (day 7), *Methylocystis* sp. strain LW5 (day 7), *Methylomonas* sp. strain LW13 (day 3). Images are representative of three independent experiments. (**C)** Methane (blue) and oxygen (black) gradients of inoculated (solid) and sterile (open) syringes after three days of incubation. Both substrates are depleted at the same depth as the horizontal band (gray bar) that forms within 25-30 mm from the filter tip.

Different strains of methanotrophs reproducibly formed the band at different rates and depths when they were individually cultured in the gradient syringe (**Fig 1B**). The gammaproteobacterium LW13 formed a 1 mm thick band within three days of incubation at a depth range within 25-30 mm from the oxygen side of the syringe. *Methylobacter tundripaludum* 21/22 formed a 0.5 mm thick band within four days of incubation at a depth range of 18-24 mm within the gradient syringe. When two methanotrophic alphaproteobacteria were individually cultured in the gradient syringe, they formed bands that were less opaque and more diffuse than the gammaproteobacteria. *Methylosinus* sp. LW3 formed a 2 mm thick band within seven days of incubation at a depth of 5-10 mm. *Methylocystis* sp. LW5 formed a band within four days at a depth of 14-19 mm and a thickness of 2 mm. Together, these findings show the ability to identify reproducible differences between methanotrophic taxa cultured in this system.

### A methane-oxygen counter gradient forms in the syringe model ecosystem

To determine if the syringe model ecosystem contained a methane-oxygen counter gradient, we measured the dissolved oxygen and methane content throughout LW13-inoculated and cell-free syringes after three days of incubation. In inoculated syringes, the dissolved oxygen concentration of the agarose closest to the filter tip was 336.9 ± 27.6 μmol/L and steeply declined to <1 μmol/L at the same depth as the horizontal band, around 25-30 mm from the filter tip (**Fig 1C**). Dissolved oxygen remained depleted with a mean concentration of 0.28 ± 0.09 μmol/L for the remainder of the agarose. This dissolved oxygen gradient began to form within 24 hours of flushing a methanotroph-inoculated gradient syringe and became steeper over the course of the three-day incubation period (**Fig S1**).

From the filter tip, the concentration of methane was <2 μmol/L until reaching the depth of the horizontal band, where it then increased in the remaining agarose up to 104.7 ± 42.67 μmol/L in the second-deepest agarose segment (**Fig 1C**). In cell-free sterile gradient syringes, a partial oxygen-methane counter gradient formed, but only reached a low of 270.6 ± 32.6 μmol/L dissolved oxygen and 5.79 ± 2.26 μmol/L methane in either gas gradient. These results indicate that bacterial consumption of methane and oxygen help drive the formation of the methane-oxygen counter gradient in this system, and that the intersection of the counter gradient is at the same location as the horizontal band.

### Horizontal band formation is correlated with increased polysaccharide content but not increased viable cell counts

We initially hypothesized that the horizontal band contained a larger number of growing cells than the surrounding depths of the syringe. To test this, we counted the colony forming units (CFUs) per mL of agarose for inoculated syringes at different depths for each methanotroph strain (**Fig 2A**). For all four methanotrophic strains tested, no individual segment contained a significantly higher CFUs/mL than any other segment from the same syringe. We confirmed this finding for LW13 using a viability stain and flow cytometry **(Fig S2)**. These results indicate that the formation of the horizontal band is not caused by increased viable cell numbers at that location.

**FIGURE 2.**
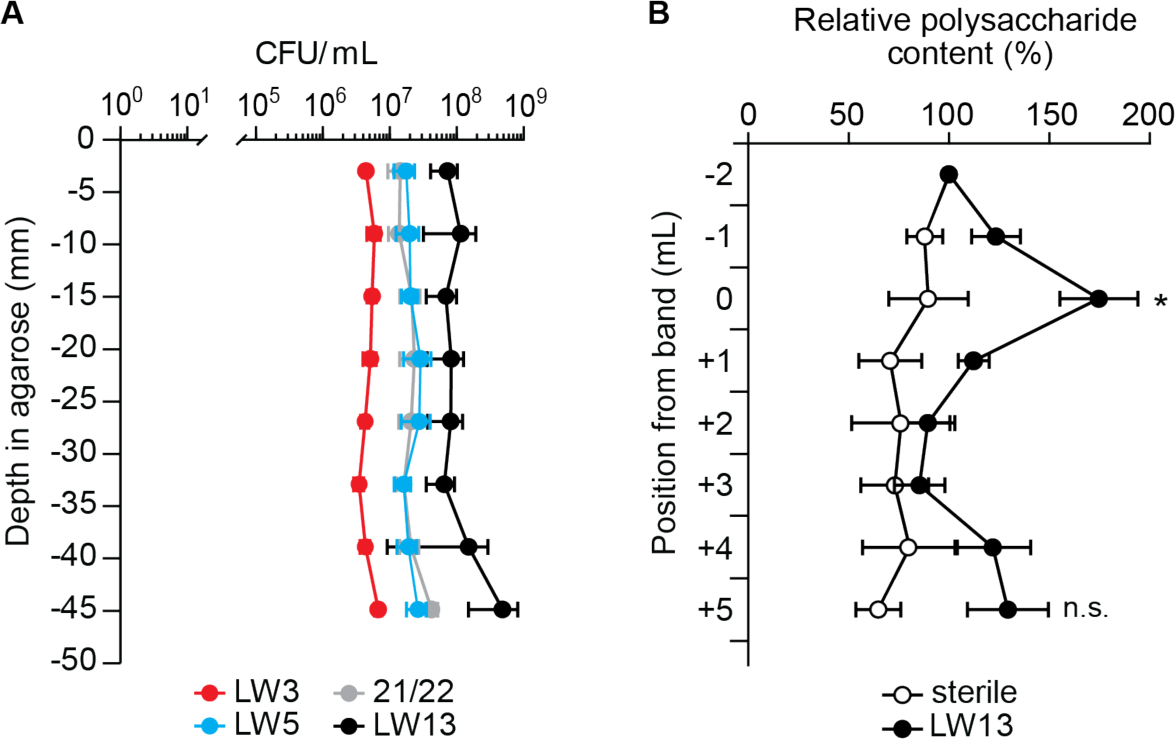
(**A)** Distribution of viable methanotrophic cells in the gradient syringe model ecosystem. The mean CFU/mL from any segment of any strain is not significantly different than any other segment mean CFU/mL of the same strain; one-way ANOVA (*p* <0.05). (**B)** Relative polysaccharide content from 1 mL agarose segments, with 0 representing a 1 mL segment centered around the horizontal band. Individual absorbance values for each syringe replicate are reported as a percentage of the absorbance at 490 nm measured at the segment closest to the filter tip from the same syringe. Data show the mean ± standard deviation of three independent experiments with two technical replicates each. *, significantly different from all other LW13 segment datapoints except the +5 point denoted n.s. (not significant); two-tailed heteroscedastic *t* test (*p* <0.05).

Because methanotrophs are known to produce polysaccharides under different growth conditions [20–22], we used a colorimetric assay [23, 24] to quantify the relative polysaccharide content at different depths in the gradient syringe. After seven days of incubating LW13 with daily methane replenishment to increase the overall opacity and thickness of the band, we divided the agarose plug into 1 mL segments, ensuring the band was centered within a single segment.

In sterile gradient syringes, only the agarose matrix contributed to the colorimetric response of the assay, and thus the relative polysaccharide content at increasing depths remained constant (**Fig 2B**). In agarose segments inoculated with LW13, the segment containing the horizontal band had significantly increased (*p* <0.05) relative polysaccharide content compared to the other segments down to 33 mm in depth. The segment furthest from the filter tip trended upwards in relative polysaccharide content, which was consistent with the observation that a faint band occasionally formed at the interface of the agarose plug and the methane-filled headspace. Because these trends correlated to the observed band position(s) within the gradient syringe, we conclude that the horizontal band formed as a result of increased polysaccharide production by cells positioned at the intersection of the methane-oxygen counter gradient.

### Polysaccharide band formation within the gradient syringe is correlated to changes in gene expression

To investigate the cellular functions underlying the spatially resolved polysaccharide-producing phenotype observed in the gradient syringe, we compared transcriptomes of LW13 isolated from increasing depths in the agarose matrix. Late exponential/early stationary phase (**Fig S3**) gradient syringes inoculated with LW13 were divided into three 2 mL segments, with the middle segment centered around the polysaccharide band. These segments were designated “above”, “band”, and “below.” Oxygen and methane profiles (**Fig 1C**) indicated differing gas substrate ratio for cells in these segments: high oxygen-low methane, low oxygen-low methane, and low oxygen-high methane, respectively. RNA-seq analysis of these segments revealed distinct gene expression profiles, with cells in the band and below segments exhibiting the greatest similarity (**Fig 3** and **S4**). Using a threshold of |log_2_(fold change)| >1.5 and adjusted *p* <0.0001, we identified 11 differentially expressed genes (DEGs) between the band and the below segments, and 312 DEGs between the band and the above segments.

**FIGURE 3.**
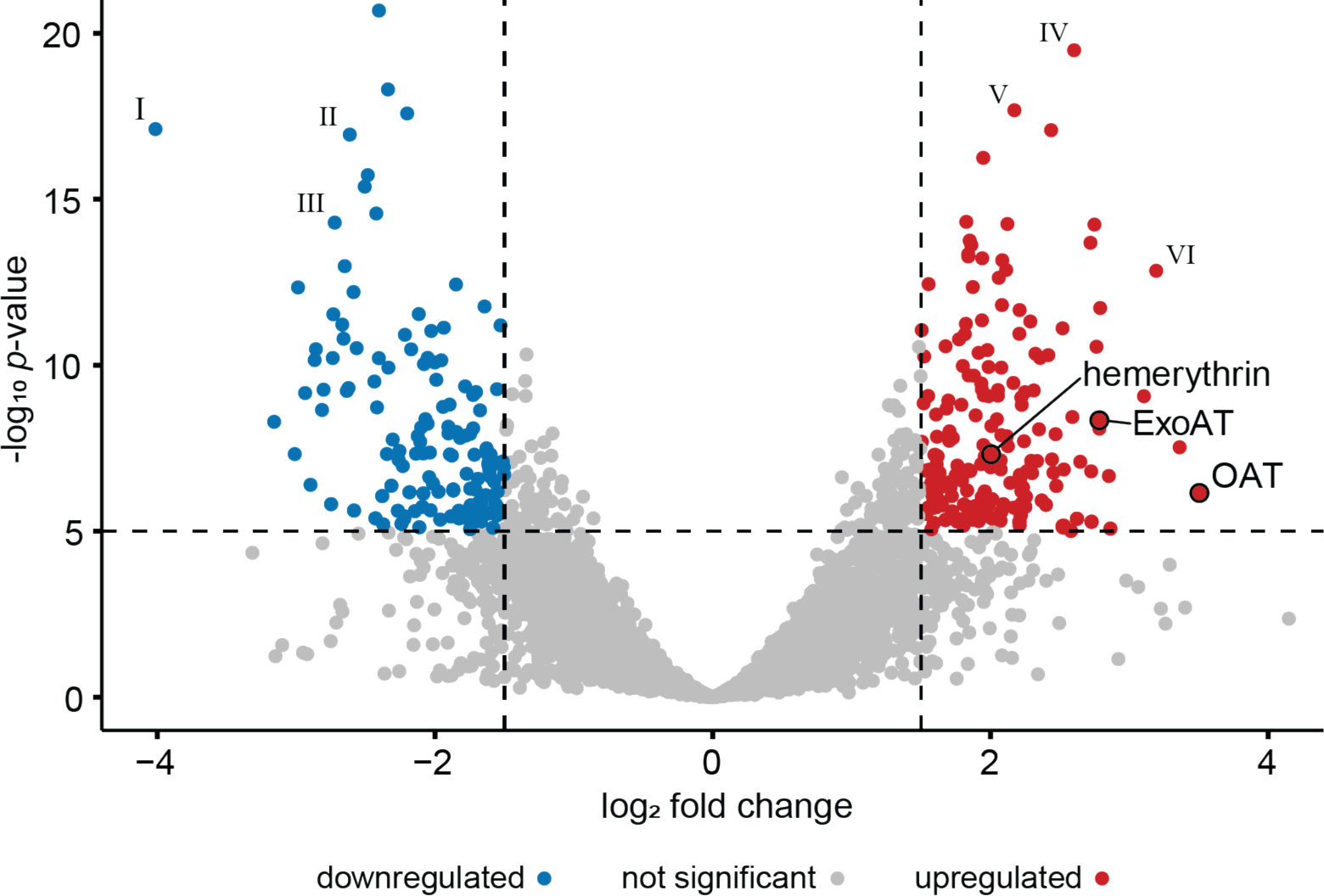
Differentially expressed genes in the band segment compared to the above segment. Dashed vertical lines signify cutoffs of |log_2_-fold change| >1.5 and the horizontal line indicates a cutoff of adjusted *p* value <0.0001. Predicted functions of highlighted genes: (I) transposase-like protein, (II) heme oxygenase, (III) polyhydroxybutyrate depolymerase, (IV) tryptophan synthase alpha chain, (V) acetyl-CoA carboxylase carboxyl transferase subunit beta, (VI) ornithine carbamoyltransferase.

In the low oxygen micro-environment of the band segment, methane metabolism genes were generally downregulated compared to the higher oxygen above segment. The *pmoCAB* methane monooxygenase operon showed an average of 1.26 log_2_-fold decrease in band cells, while the *mxaF* and *xoxF* methanol dehydrogenase genes exhibited decreases of 0.623 and 1.02 log_2_-fold change, respectively. Genes predicted to be involved in nitrogen transport were upregulated in the band segment. Using less stringent cutoffs, only one DEG was upregulated in the band segment compared to both above and below segments: a hypothetical protein within a predicted Type VI secretion system gene cluster. In comparing gene expression between band and below segments (**Fig S4A**), there were notably fewer DEGs with a log_2_-fold change >1.5 than in the band vs. above comparison, presumably due to the shared low oxygen micro-environments of the band and below segments. Among 11 upregulated DEGs were two genes predicted to be involved in iron acquisition and storage, and three genes predicted to be involved in cell division.

Comparing the gene expression between cells from the band segment and those from the above segment revealed a large range of DEGs with a log_2_-fold change >1.5, highlighting potential differences in cellular activity in these segments. In the band segment cells, there was significant upregulation of genes annotated as involved in energy production, amino acid and nucleotide metabolism/transport, coenzyme metabolism, and carbohydrate and lipid metabolism/transport compared to the above segment, as assigned by the NCBI Cluster of Orthologous Genes database [25]. These genes made up 78 of the 177 upregulated genes, suggesting involvement in diverse metabolic activities including energy conservation and synthesis of biomolecules. Additionally, 15 upregulated genes were predicted to be related to cellular structure and motility. Collectively, these upregulated genes suggest intensified activity in metabolism, biosynthesis, translation, transcription, cellular structure, and intracellular processes. Moreover, the increased expression of 17 genes predicted to be involved in intracellular trafficking and secretion, signal transduction, and secondary structure suggests enhanced intracellular communication and responsiveness to external conditions. Using less strict cutoffs, we found several putative functional pathways that were also upregulated, including electron transport, type VI secretion, hopanoid biosynthesis, and cell wall biosynthesis. These results reveal differences in gene expression within this spatially resolved model ecosystem.

### Differentially expressed genes are involved in polysaccharide band formation

Among the upregulated genes with the greatest log_2_-fold change in the band compared to the above segment, we selected three for further investigation. We chose genes with annotated functions that to our knowledge had not been extensively studied in methanotrophs, especially in the context of polysaccharide production. These genes were predicted to encode a fucose 4-O-acetyltransferase (OAT; gene ID: 2923715777, Integrated Microbial Genomes System [26]), a hemerythrin (2923717515), and an *N*-acyl amino acid synthase of a PEP-CTERM/exosortase system (ExoAT) (2923716464). We constructed disruption mutant strains for each of the three genes to determine their individual influence on the growth of LW13 and the formation of the polysaccharide band within the gradient syringe (**Fig 3**). We disrupted the target genes via electroporation to create the mutant strains ΔOAT, Δhemerythrin, and ΔExoAT [27]. When we individually inoculated these LW13 mutants into gradient syringes, we observed the mutant strains did not form the same distinct polysaccharide band, and that band formation was restored after gene complementation (**Fig 4A**). We also counted the viable cells in each syringe after six days of incubation with daily methane replenishment to determine if these genes were involved in cell growth within the system. While wild-type LW13 grew to more than 75-times its initial CFU/mL concentration in the gradient syringe, the number of viable LW13ΔOAT, LW13Δhemerythrin, and LW13ΔExoAT mutant cells did not significantly (*p* <0.05) increase over the growth period (**Fig 4B**). Complementation of the deleted genes led to significantly increased growth in the three mutants from Day 0 to Day 6, restoring final CFU/mL to values comparable to the LW13 wild-type strain. Deletion of these genes did not negatively affect growth in planktonic culture, but instead led to slight but significantly faster growth rates (**Fig S5**). Together, these results show we can use the model gradient syringe system to link genes to phenotypes that would otherwise be missed using conventional culturing methods.

**FIGURE 4.**
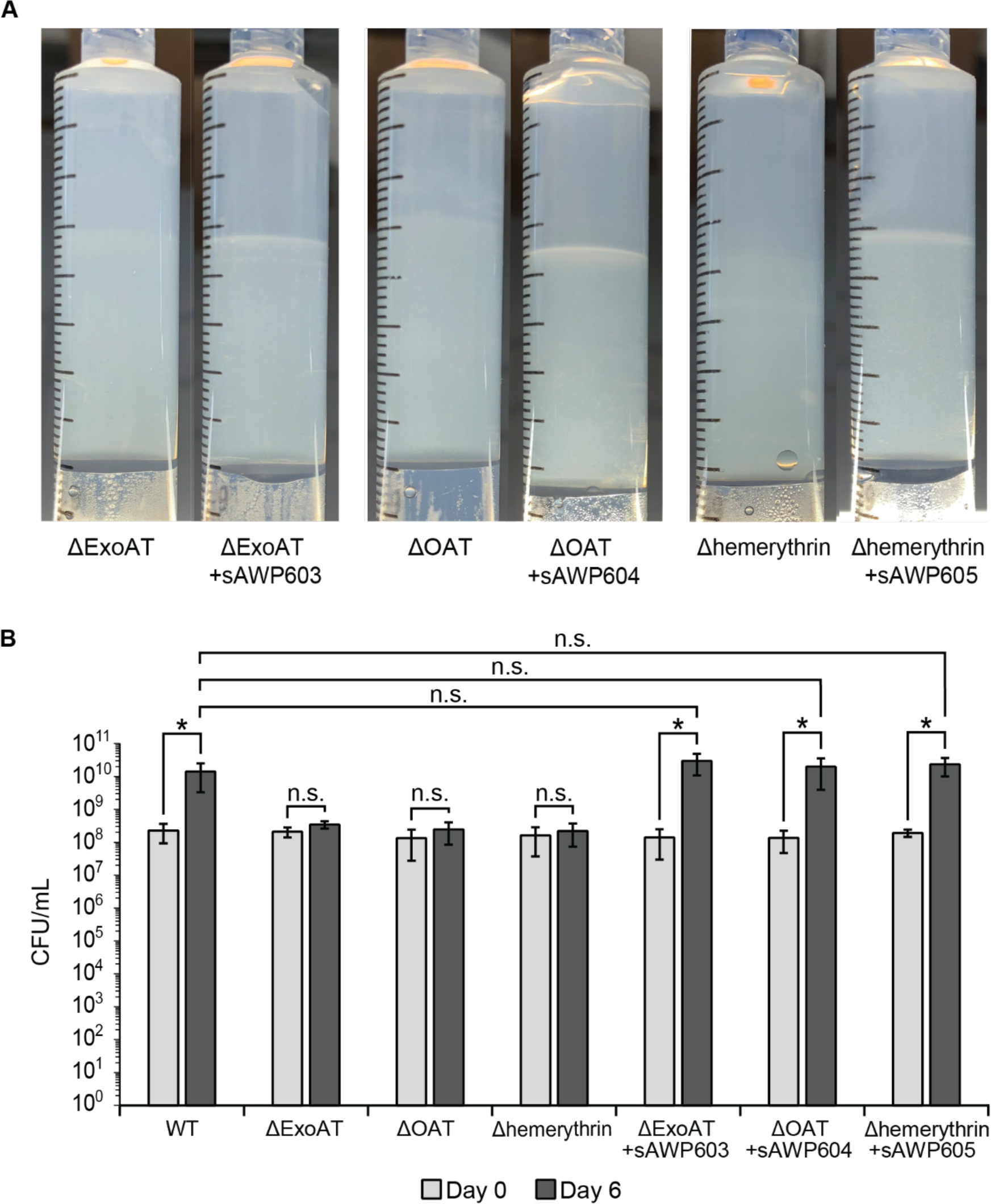
(**A)** Reduction in polysaccharide band production in three LW13 mutants within the gradient syringe. Images are representative of three independent experiments. (**B)** Growth of LW13 strains in the gradient syringe over a six-day incubation period. Data show the mean ± standard deviation of three independent experiments with two technical replicates each. *, significantly different (two-tailed heteroscedastic *t* test, α = 0.05). n.s., not significantly different.

## DISCUSSION

Methanotrophs use oxygen to catabolize methane as their sole carbon and energy source and often exist within a counter gradient of these gaseous substrates in the environment. In this work we observed that methanotrophs form a distinct horizontal band when grown in a spatially resolved model ecosystem containing a methane-oxygen counter gradient. We determined that the band is located at the intersection of the counter gradient and that it is not due to increased numbers of viable cells at this location, as previously hypothesized [16–18], but instead due to increased polysaccharide content at this location. We then used transcriptomics and deletion mutants to identify three genes involved in this phenotype. These genes are also required for cell growth within the model ecosystem, but their absence did not negatively affect the ability of LW13 to grow in planktonic culture.

The observed increase in polysaccharide content in the horizontal band in the gradient syringe model ecosystem may be due to increased extracellular polymeric substances (EPS) produced by methanotrophs at this depth. An established link exists between methane oxidation, oxygen availability, and EPS production in methylotrophic bacteria [20, 28]. However, more experiments will be necessary to determine the exact molecular components of this band, as well as whether the polysaccharides are within or outside of methanotrophic cells. Previously, the role of the exosortase protein EpsH in production of EPS in LW13 was identified [29] and confirmed [30], where disruption of *epsH* resulted in significantly reduced EPS in ammonium mineral salts media. However, this gene was not significantly differentially expressed by cells in any segment in our RNA-seq analysis (**Fig 3** and **S4**), possibly due to nitrate, rather than ammonium, being used as the nitrogen source in our experiments.

We observed variations in the depth and appearance of the horizontal band when we grew different methanotrophic genera in the gradient syringe (**Fig 1B**). This may stem from different methanotrophic genera being adapted to different oxygen tensions the environment [10, 11, 31]. It is unclear whether formation of the polysaccharide band is a cause or consequence of this location being at the intersection of the methane-oxygen counter gradient. It is possible that polysaccharide production is higher in cells at the intersection of the counter gradient as a response to low gas concentrations at this location, or that polysaccharide production led to higher gas consumption rates at this depth. While the exact triggers for polysaccharide production in methanotrophs remain to be precisely defined, excess carbon, nitrogen deficiency, and oxygen excess or limitation have all been reported to stimulate EPS production in methylotrophs [20, 21, 32]. Determining these factors may have practical implications for applied methanotrophy, such as methane mitigation in landfill covers and biofilters, which are negatively affected by the production of EPS [28].

Transcriptomic analysis of different segments within an LW13-inoculated gradient syringe revealed broad global gene expression differences, with band and below segments being most similar (**Fig S3A**), suggesting that oxygen availability is a major influence on cellular heterogeneity. To explore possible gene-phenotype connections, we investigated a subset of genes that were upregulated in the band segment that had not yet been experimentally linked to polysaccharide production in methanotrophs. *N*-acyl amino acid synthases have been phylogenetically linked to the PEP-CTERM/exosortase system (ExoAT) [33], a putative protein sorting system in Gram-negative bacteria associated with EPS production [34]. The predicted product of the fucose 4-O-acetylase like acetyltransferase gene (OAT) likely engages in acylation processes impacting various sugar-containing cell-surface structures, including exopolysaccharides [35]. Hemerythrin is a non-heme oxygen-binding protein that serves as an oxygen scavenging transporter for pMMO-containing aerobic methanotrophs such as LW13 [31]. Single knockouts of these genes in LW13 had a negative effect on horizontal band formation (**Fig 4A**) and growth (**Fig 4A**) in the gradient syringe model ecosystem, but did not interfere with growth in planktonic culture (**Fig S5**). The fact that band formation and growth were both affected in each of the mutants means that it is possible that methanotrophic growth is dependent on horizontal band formation in this system, or that formation of the band is dependent on growth.

In this work we demonstrate that culturing methanotrophs within a spatially resolved counter gradient of methane and oxygen can reveal gene-phenotype associations that are largely unnoticed in conventional culturing approaches. Our results have raised questions about methanotroph physiology within a methane-oxygen counter gradient that can now be studied at the molecular level using this model ecosystem. More broadly, this study highlights the utility of even simple laboratory-based model ecosystems for linking genes with organismal phenotypes.

## ACKNOWLEDGEMENTS

This work was supported by startup funding from the University of Utah Department of Chemistry. We thank members of the Puri Lab for helpful discussions. We thank Rachel Hurrell (University of Utah) for initial guidance with the flow cytometry experiment.

## AUTHOR CONTRIBUTIONS

D.G.B. and A.W.P. designed the experiments, analyzed the data, and wrote the manuscript. D.G.B. performed the experiments. D.G.B. and A.W.P. edited and approved of the final version of the manuscript.

## DATA AVAILABILITY

The datasets generated and analyzed during the current study are available in the NIH NCBI Gene Expression Omnibus (GEO) database under accession number GSE243827. This includes raw transcriptomic data, as well as normalized counts and computed pairwise fold changes for the different syringe segments.

## METHODS

### Strain growth

Liquid cultures of *Methylomonas* sp. LW13 were grown in an atmosphere of 50% methane in air in liquid nitrate mineral salts (NMS) medium containing 0.2 g/liter MgSO_4_·7H_2_O, 0.2 g/liter CaCl_2_·6H_2_O, 1 g/liter KNO_3_, and 30 μM LaCl_3_ as well as 1X trace elements; 500X trace elements contain 1.0 g/liter Na_2_-EDTA, 2.0 g/liter FeSO_4_·7H_2_O, 0.8 g/liter ZnSO_4_·7H_2_O, 0.03 g/liter MnCl_2_·4H_2_O, 0.03 g/liter H_3_BO3, 0.2 g/liter CoCl_2_·6H_2_O, 0.6 g/liter CuCl_2_·2H_2_O, 0.02 g/liter NiCl_2_·6H_2_O, and 0.05 g/liter Na_2_MoO·2H_2_O. A final concentration of 5.8 mM phosphate buffer (pH 6.8) was added before use. Liquid cultures were grown in 18 x 150 mm tubes sealed with rubber stoppers and aluminum seals, and were shaken at 200 rpm at room temperature.

### Preparation of gradient syringes

Liquid cultures of methanotrophs in exponential-phase growth (OD_600_ ∼ 0.5) were used to inoculate gradient syringes. Cells were pelleted and resuspended in sterile water to reach an approximate OD_600_ of 1.0. For each 10 mL syringe, 1 mL of concentrated cells was mixed with 5 mL NMS1 and 4 mL molten agarose (0.5% w/v, cooled to 55°C). The mixture was slowly poured to the 8 mL marking of a 10 mL plastic Luer-Lok tip syringe (BD) fitted with a sterile PTFE filter tip (Titan3 PTFE (Hydrophobic) Syringe Filter 0.2 µm, 4 mm, Thermo Scientific). Solidified syringes were capped with a sterile 20 mm rubber butyl stopper (Chemglass Life Sciences) and wrapped with lab tape for labeling. Inlet and outlet needles (23 gauge, BD) were pierced through the rubber stopper, and 20 mL of 100% CH_4_ was flushed through the syringe headspace. Syringes were flushed daily with methane and incubated with the PTFE filter pointing up at 18°C.

### Gas analysis of gradient syringes

The dissolved oxygen within the gradient syringe was measured using a Clark-type microelectrode (Ox50, Unisense) at 0.1 mm intervals. Immediately before measurement, the tip of a syringe was sliced off with a razor and the syringe fastened to a syringe pump (NE-1000, New Era Pump Systems Inc.) that moved the agarose plug towards the microelectrode at a rate of 1 mL per minute (0.6 cm/min). Measurements were recorded and converted to μmol/L using the Unisense Logger software. The relative methane concentration in LW13-containing and cell-free gradient syringe agarose were analyzed using gas chromatograph-flame ionization detector. Just before extrusion, the syringe filter was replaced with a two-way stopcock with a 23-gauge needle and the syringe’s rubber stopper was quickly exchanged with a syringe plunger. Eight 1 mL agarose aliquots were then added to different evacuated 12 mL Exetainer vials (Labco). Prior to injecting 500 µL of headspace into an Agilent 6890N gas chromatograph, sample vials were equilibrated to atmospheric pressure. A calibration curve employing known CH_4_ amounts in vials was used to convert peak area (pA*min) to μmol/L.

### Counting of viable methanotroph cells in gradient syringe

Within one day of horizontal band formation, agarose from the gradient syringe was extruded in 1 mL aliquots through a 23-gauge sterile needle using the accompanying syringe plunger. Agarose was diluted with 1 mL sterile water and vortexed to homogenize the samples and aid in pipetting. Samples were serially diluted tenfold in sterile water before spotting onto solid NMS agar plates. Plates were incubated in sealed jars under 40% methane in air and grown at 18°C. Bacterial colonies were counted and used to calculate the colony forming units per milliliter (CFUs/mL).

For flow cytometry of LW13-containing syringes, after 7 days of growth, each 1 mL aliquot of extruded agarose was diluted and homogenized with the addition of 0.75 mL of 0.85% NaCl in water. Aliquots were further diluted 1:10 with the salt solution before staining 1:1 with a mixture of SYTO9 and propidium iodide (LIVE/DEAD BacLight Bacterial Viability Kit, Invitrogen) and incubating in the dark at room temperature for 15 minutes. Samples were analyzed with a CytoFLEX flow cytometer (Beckman Coulter) with parameters: 30000 events; fluidics: slow; excitation: 470 nm; emission: 490-700 nm. Stained aliquots from cell-free syringes and 70% ethanol-treated aliquots were used as negative controls to determine voltage gates. Live cell events per μL values were used to compare viable cell concentrations of different agarose segments.

### Polysaccharide detection assay

One milliliter aliquots of agarose were extruded into 2 mL microcentrifuge tubes and mixed with 1 mL of a 1% (m/v) Na_2_CO_3_ solution in water. Samples were heated to 80°C for 30 minutes and vortexed every 5-10 minutes, followed by centrifugation at 4,000 x g at 4°C for 20 minutes. The supernatant was collected and combined with three volumes of 100% ethanol and incubated at −20°C overnight. The ethanol-precipitated polysaccharides were collected by centrifugation at 16,100 x g at 4°C for 30 minutes, and the air-dried pellet was resuspended in 100 uL of Milli-Q water. The relative polysaccharide content of each agarose segment was measured using a phenol-sulfuric acid colorimetric assay. Briefly, 50 μL of resuspended extract was combined with 150 μL concentrated sulfuric acid and 30 uL of 5% phenol in a clear 96-well plate [20, 23, 24]. The absorbance at 490 nm was measured using a microplate reader, and the relative polysaccharide content of individual agarose segments was calculated as a percentage of the absorbance at the agarose segment closest to the PTFE filter.

### RNA preparation, RNA-seq, and data analysis

For each of three independent replicates, eight *Methylomonas* sp. LW13-inoculated syringes were incubated for three days to reach exponential growth. Agarose segments (2 mL) positioned above, at, and below the polysaccharide band were extruded from each syringe through a 23-gauge needle into microcentrifuge tubes. Samples were frozen at −80°C until RNA extraction. Thawed aliquots were centrifuged at 21,000 x g at 4°C for 15 minutes. After removal of the supernatant, 600 µL of pre-warmed CTAB extraction buffer was added. The CTAB buffer consisted of 2% CTAB, 2% polyvinylpyrrolidone 40, 2 M sodium chloride, 100 mM Tris-HCl (pH 8.0), and 20 mM EDTA [36]. Zirconia glass beads were added, and samples were homogenized for 3 minutes at 30 Hz/sec using a bead beater. Samples were extracted twice with 600 µL chloroform:isoamyl alcohol (24:1), and the aqueous phase was mixed with isopropanol and incubated overnight at −20°C. Precipitates were centrifuged for 30 minutes at 16,100 x g at 4°C for 30 minutes and pellets were washed twice with cold 75% ethanol made with DEPC water. Air dried pellets were dissolved in 100 uL DEPC water and treated with DNase I (Ambion) at 37°C for 30 minutes. The DNase I was inactivated by extraction with three volumes of acid phenol:chloroform:IAA (125:24:1, pH 4.5) before pooling RNA from the same segment across all syringes in an overnight precipitation at −20°C in 1 volume isopropanol and 0.8 M LiCl. Precipitates were washed twice with cold 70% ethanol made with DEPC water, air dried, and resuspended in 50 uL DEPC water. Samples were re-purified using RNA Clean & Concentrator-5 (Zymo Research) to remove small RNAs.

Library preparation and sequencing were performed by the High-Throughput Genomics Shared Resource Core at the University of Utah. cDNA libraries were prepared using a NEBNext Ultra II Directional RNA Library Prep Kit with rRNA Depletion (New England Biolabs) and sequenced using an Illumina NovaSeq 6000 with 150×150 cycle paired end sequence run. RNA sequencing data analysis was conducted using a KBASE workflow [37]. Reads quality was assessed using FastQC v0.11.9, trimmed with Trimmomatic v0.36 [38], aligned using HISAT2 v2.1.0 [39], transcripts assembled with StringTie v2.1.5 [40], and differentially expressed genes identified using DESeq2 v1.20.0 [41].

### Genetic manipulation in *Methylomonas* sp. LW13

Plasmids and constructs used in this study are listed in **Table S1**, and primers are listed in **Table S2**. All genetic manipulation of LW13 was carried out at 30°C under 50% methane in air.

#### Deletion mutant construction

Genetic manipulations were accomplished by electroporating PCR amplification-generated constructs into LW13, followed by the selection of chromosomal recombinants, following established methods [27]. Briefly, the kanamycin resistance gene cassette (amplified from plasmid pCM433kanT [42]) and flanking regions of the target gene were assembled using overlap extension PCR. The assembled fragment was then amplified using nested primers and purified by gel purification. After confirming correct assembly via Sanger sequencing the fragment was electroporated directly into LW13 cells. Post-electroporation, cells were cultured overnight in NMS1 medium without antibiotics, and then transferred to kanamycin-containing plates (50 µg/ml). Single colonies were selected, and disruption of the gene of interest with the kanamycin resistance cassette was confirmed through colony PCR using OneTaq with primers specific to the assembled construct.

#### Complementation of genes into mutant strains

Using overlap extension PCR, ∼3000-nucleotide sequences containing the gene of interest were constructed and electroporated into LW13 mutants. These constructs included the gene of interest under the bacterial promoter *nptII* [43], along with a zeocin resistance gene cassette directly downstream of the gene to be complemented. Flanking either end of this construct were homologous flanks targeting an insertion site in the LW13 genome that was previously used for complementation in this strain [44]. Full-length DNA segments were excised after separation on a 1% TAE agarose gel and purified using a ThermoFisher GeneJet Gel Purification Kit. After introduction of the segments via electroporation, successful recombinants were selected on NMS1 media containing 20 μg/mL zeocin and confirmed by colony PCR.

