## Supplementary material for "Methanotroph phenotypic heterogeneity in a methane-oxygen counter gradient": Figs S1-S5; Tables S1-S2

**SUPPLEMENTARY FIGURES**

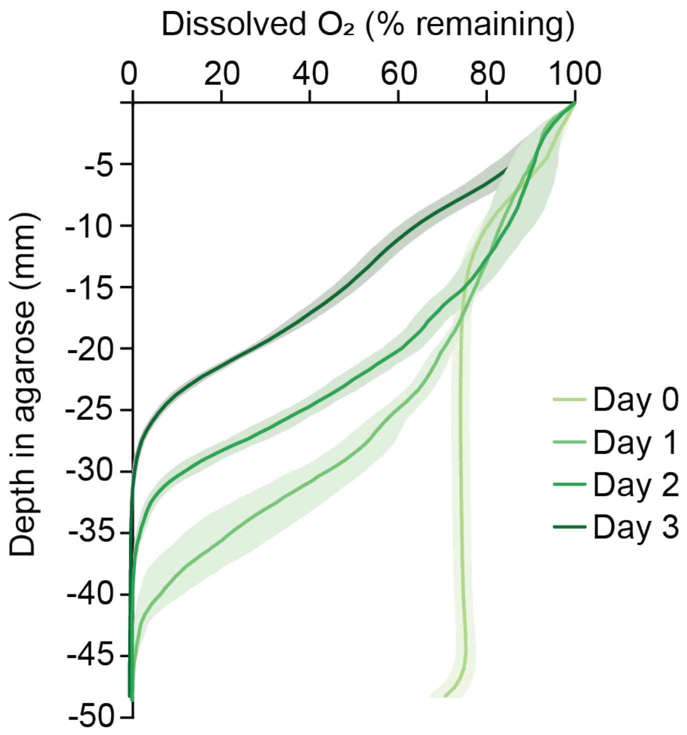


**Figure S1.** Development of the oxygen gradient in LW13-inoculated gradient syringes over three days. Data show the mean ± standard deviation (shaded areas) of three independent experiments with five technical replicates each.

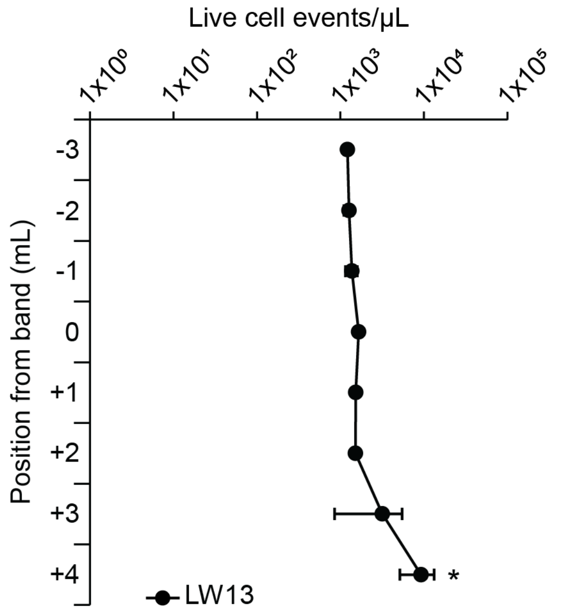


**Figure S2.** Distribution of cells in LW13-inoculated syringes incubated for 7 days. Live cell events per µL of diluted agarose were quantified by flow cytometry. Cell concentrations in the deepest segment +4 mL from the band were significantly higher than all other segments (one-way ANOVA with Tukey-Kramer post hoc analysis, $p < 0.05$). Data points and values represent the mean ± standard deviation of two independent experiments with two technical replicates each.

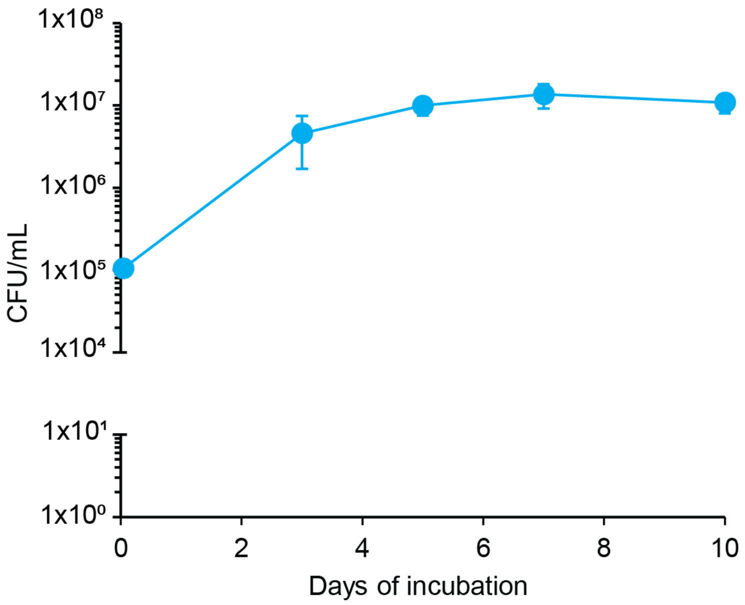


**Figure S3.** Viable cell counts of LW13-inoculated gradient syringes over ten days. Data show the mean ± standard deviation of three independent experiments with two technical replicates each.

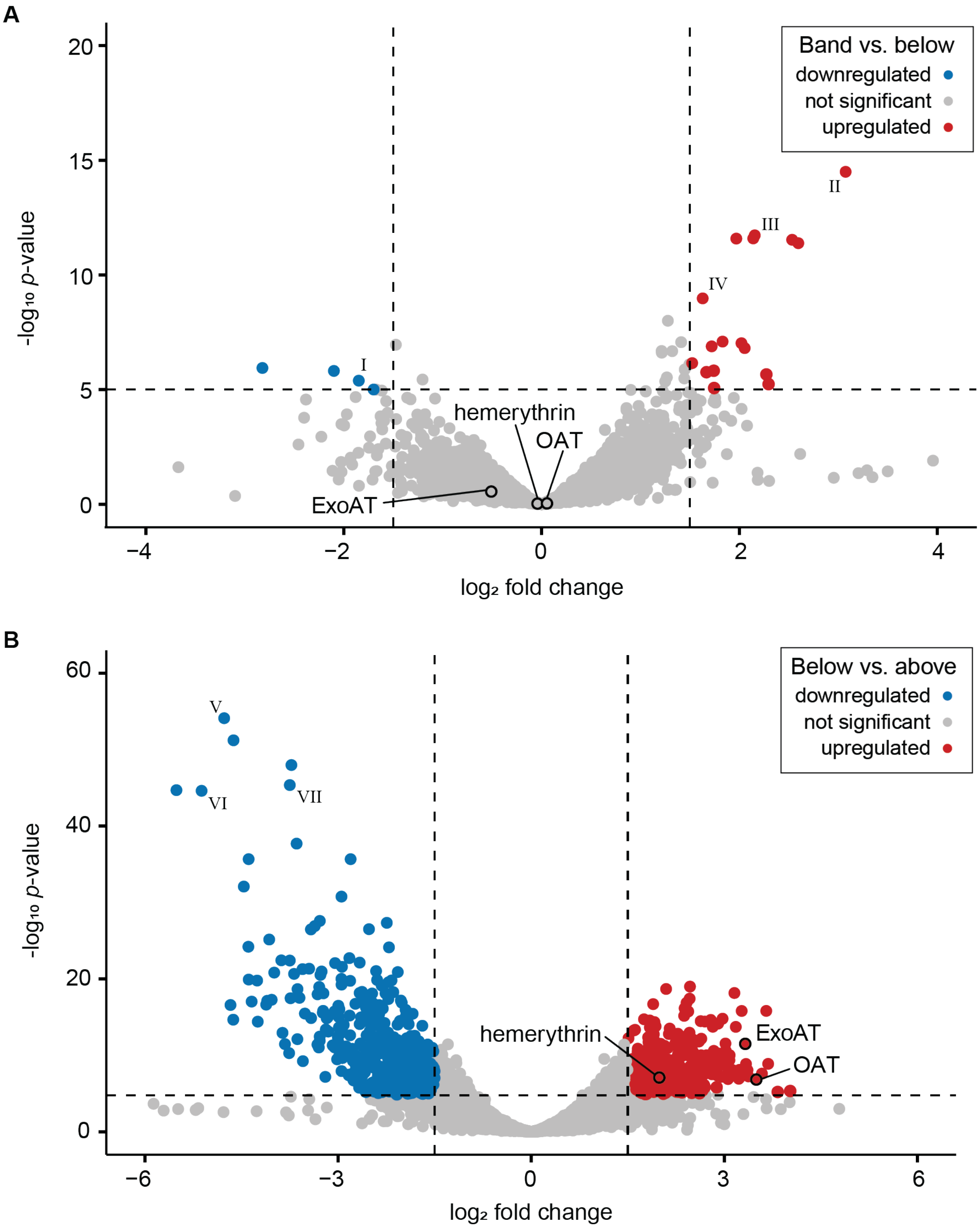


**Figure S4.** Differentially expressed genes comparing different gradient syringe segments. Dashed vertical lines signify cutoffs of $|\log_2\text{-fold change}| > 1.5$ and the horizontal line indicates a cutoff of adjusted *p* value <0.0001. **A** Band vs. below comparison with highlighted significantly differentially expressed genes (I) bacterioferritin-associated ferredoxin, (II) heme oxygenase, (III) cell division protein FtsA. **B** Below vs. above comparison with highlighted significantly differentially expressed genes (IV) heme oxygenase, (V) bacterioferritin-associated ferredoxin, (VI) two-component response regulator RegA.

**A**

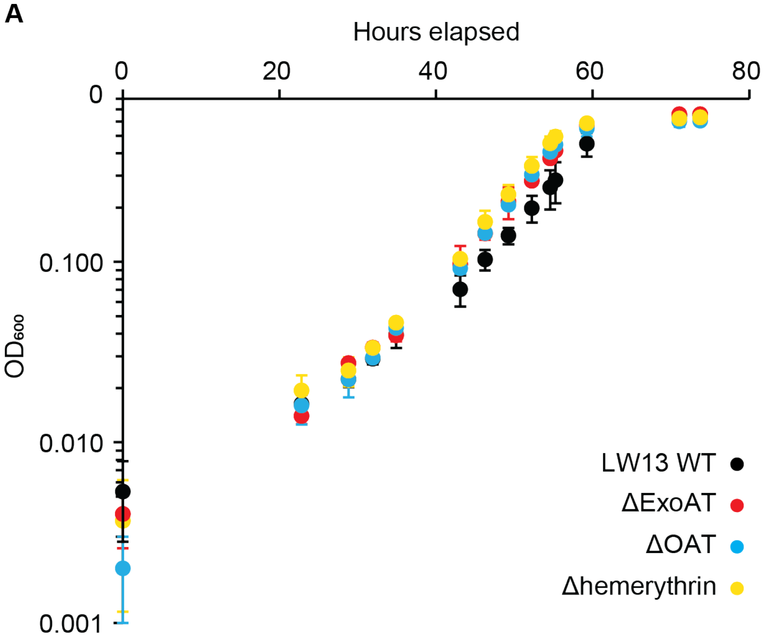


**B**

| strain | doubling time (h) |
|---|---|
| LW13 WT | 6.81 ± 0.34 |
| ΔExoAT | 6.13 ± 0.11 * |
| ΔOAT | 6.19 ± 0.19 * |
| Δhemerythrin | 6.20 ± 0.17 * |

**Figure S5.** **A** Growth curve of wild-type LW13 and single deletion mutants in shaken liquid nitrate mineral salts (NMS) media at room temperature. **B** Doubling times in hours of strains in liquid culture; mutant doubling times were significantly lower than the wild-type control (one-way ANOVA with Tukey-Kramer post hoc analysis, $p < 0.05$). Data points and values represent the mean ± standard deviation of three technical replicates and are representative of three independent experiments.

### SUPPLEMENTARY TABLES

**Table S1.** Strains and SOE constructs used in this study.

| Strain or construct | Description | Source or reference |
|---|---|---|
| **Strains** | | |
| *Methylomonas* sp. strain LW13 | Aerobic methane-oxidizing bacterium isolated from Lake Washington sediment | [1] |
| *Methylobacter tundripaludum* strain 21/22 | Aerobic methane-oxidizing bacterium isolated from Lake Washington sediment | [1] |
| *Methylosinus* sp. LW3 | Aerobic methane-oxidizing bacterium isolated from Lake Washington sediment | [1] |
| *Methylocystis* sp. LW5 | Aerobic methane-oxidizing bacterium isolated from Lake Washington sediment | [1] |
| *Methylomonas* sp. strain LW13ΔOAT | Strain containing deletion of gene 2923715777 with predicted fucose 4-O-acetylase-like acetyltransferase product | Locus tag: Ga0485366_01_330073_331065 (https://img.jgi.doe.gov) |
| *Methylomonas* sp. strain LW13Δhemerythrin | Strain containing deletion of gene 923717515 with predicted hemerythrin product | Locus tag: Ga0485366_01_2188896_2189291 (https://img.jgi.doe.gov) |
| *Methylomonas* sp. strain LW13ΔExoAT | Strain containing deletion of gene 2923716464 with predicted N-acyl amino acid synthase of PEP-TERM/exosortase system product | Ga0485366_01_1079880_1080629 (https://img.jgi.doe.gov) |
| **Construct** | | |
| sAWP507 | SOE construct to create LW13ΔOAT; kanR cassette flanked by ~800 bp regions directly adjacent to 2923715777 | This study |
| sAWP508 | SOE construct to create LW13Δhemerythrin; kanR cassette flanked by ~800 bp regions directly adjacent to 2923717515 | This study |
| sAWP510 | SOE construct to create LW13ΔExoAT; kanR cassette flanked by ~800 bp regions directly adjacent to 2923716464 | This study |
| sAWP603 | SOE construct to complement LW13ΔExoAT with 2923716464 under *nptII* promoter between 2923716887 and 2923716888 in genome | This study |
| sAWP604 | SOE construct to complement LW13ΔOAT with 2923715777 under *nptII* promoter between 2923716887 and 2923716888 in genome | This study |
| sAWP605 | SOE construct to complement LW13Δhemerythin with 2923717515 under *nptII* promoter between 2923716887 and 2923716888 in genome | This study |

**Table S2.** Cloning primers used in this study.

| Primer name | Sequence (5' to 3')* | Description |
|---|---|---|
| **Gene deletion constructs** | | |
| oAWP1238 | GATGAGAGCTTTGTTGTAGG | For amplification of kanR cassette for insertion between flanks to replace native gene |
| oAWP1239 | TCTCGAGTCCCGTCAAGTC | |
| oAWP1646_507U_fwd | AGACGACATCGAATTTAAACTTCGC | For amplification of flanks to construct sAWP507 to create LW13ΔOAT |
| oAWP1647_507U_rev | AACTGGTCCACCTACAACAAAGCTCTCATCGCCCCAATACAACAAGGAAAATTGCA | |
| oAWP1648_507D_fwd | AGCATTACGCTGACTTGACGGGACTCGAGATGTGCCATTATTGAACTCTGTGACA | |
| oAWP1649_507D_rev | GCACACCTATCACGCCATTGCT | |
| oAWP1650_508U_fwd | GCAAAACCTGGGCGACGATTGC | For amplification of flanks to construct sAWP508 to create LW13Δhemerythrin |
| oAWP1651_508U_rev | AACTGGTCCACCTACAACAAAGCTCTCATCGGTTTGGTGCTCTTGGTCGGCA | |
| oAWP1652_508D_fwd | AGCATTACGCTGACTTGACGGGACTCGAGATCGACATGGCGTATTCTGAAGCA | |
| oAWP1653_508D_rev | AATTGGGCACGTTACGCGGGTC | |
| oAWP1658_510U_fwd | GCGGCGATGCCTCTGTGTTTCA | For amplification of flanks to construct sAWP510 to create LW13ΔExoAT |
| oAWP1659_510U_rev | AACTGGTCCACCTACAACAAAGCTCTCATCTGCGTTTGCTTTCAGGCGTGTCA | |
| oAWP1660_510D_fwd | AGCATTACGCTGACTTGACGGGACTCGAGAGAGCAATCGGTGGCTAAGCGGC | |
| oAWP1661_510D_rev | ACCACACGGCAAGCGCTAAAGC | |
| **Gene complementation constructs** | | |
| oAWP1877_2887_fwd | ACCATTAACGGCGAAGTCAGCA | for amplification of 2923716888 from LW13 genome; overlaps with *nptII* promoter |
| oAWP1927_2887_pnpt_rev | TCCCCAATTCCTGGCAGTTTATGGGTCAATCCGAGTACCACTACAGAGCT | |
| oAWP1928_pnpt_2887_fwd | ATTAGTTGTAAGCTCTGTAGTGGTACTCGGATTGACCCATAAACTGCCAG | for amplification of *nptII* promoter; 30 nt overlap with 2923716888 |
| oAWP1929_pnpt_exo_rev | ACTATCAAATGAATGCTTTTCAGAAATCAATTTTTCTTCCTCCACTAGTA | for amplification of *nptII* promoter; 30 nt overlap with ExoAT |
| oAWP1931_pnpt_fuc_rev | TATATCAACGAGTCTGTTTTTTTTATCCAATTTTTCTTCCTCCACTAGTA | for amplification of *nptII* promoter; 30 nt overlap with OAT |
| oAWP1933_pnpt_heme_rev | TTGAGCCGCAGTCCAAGTAATTAAAGCCATTTTTTTCTTCCTCCACTAGTA | for amplification of *nptII* promoter; 30 nt overlap with hemerythrin |
| oAWP1930_exo_pnpt_fwd | AGAGACAGGATACTAGTGGAGGAAGAAAAATTGATTTCTGAAAAGCATTC | for amplification of ExoAT; 30 nt overlap with *nptII* promoter |
| oAWP1888_589_zeo_rev | TGGCCATAGCTGTTTCCTGTGTGAATACCTTTAAGCTCTTTGCCGCTTAG | for amplification of ExoAT; 30 nt overlap with zeoR insert |
| oAWP1932_fuc_pnpt_fwd | AGAGACAGGATACTAGTGGAGGAAGAAAAATTGGATAAAAAAAACAGACT | for amplification of OAT; 30 nt overlap with nptII promoter |
| oAWP1890_590_zeo_rev | TGGCCATAGCTGTTTCCTGTGTGAATACCTCTAATTTGTCACAGAGTTCAATAATG | for amplification of OAT; 30 nt overlap with zeoR insert |

| | | |
|---|---|---|
| oAWP1934_heme_pnpt_fwd | AGAGACAGGATACTAGTGGAGGAAGAAAAAATGGCTTTAATTACTTGGAC | for amplification of hemerythrin; 30 nt overlap with *nptII* promoter |
| oAWP1892_591_zeo_rev | TGGCCATAGCTGTTTCCTGTGTGAATACCTTTACTTCAATGCTTCAGAATACGC | for amplification of hemerythrin; 30 nt overlap with zeoR insert |
| oAWP1883_zeo_589_fwd | CAATCGGTGGCTAAGCGGCAAAGAGCTTAAAGGTATTCACACAGGAAACA | for amplification of zeoR; 30 nt overlap with ExoAT |
| oAWP1884_zeo_590_fwd | GTGCCATTATTGAACTCTGTGACAAATTAGAGGTATTCACACAGGAAACA | for amplification of zeoR; 30 nt overlap with OAT |
| oAWP1885_zeo_591_fwd | GACATGGCGTATTCTGAAGCATTGAAGTAAAGGTATTCACACAGGAAACA | for amplification of zeoR; 30 nt overlap with hemerythrin |
| oAWP1886_zeo_2888_rev | CAGGCAAAAAAAAGCCCGCTGTATCGCGTATTATTCAGTCCTGCTCCTCG | for amplification of zeoR; 30 nt overlap with 2923716887 |
| oAWP1881_2888_zeo_fwd | ACTTCGTGGCCGAGGAGCAGGACTGAATAATACGCGATACAGCGGGCTTT | for amplification of 2923716887 from LW13 genome; overlaps with zeoR |
| oAWP1882_2888_rev | AAATTTTTACCCAAACTGCTTGGTCTC | |

*Homology regions used for SOE PCR are underlined.
